# ATP and Substrate Binding Regulates Conformational Changes of Human Peroxisomal ABC Transporter ALDP

**DOI:** 10.1101/2021.10.14.464310

**Authors:** Chao Xiong, Li-Na Jia, Ming-He Shen, Wei-Xi Xiong, Liu-Lin Xiong, Ting-Hua Wang, Dong Zhou, Zheng Liu, Lin Tang

## Abstract

The malfunction of ABCD1 causes X-linked adrenoleukodystrophy (X-ALD), a rare neurodegenerative disease that affect all tissues in human. Residing in the peroxisome membrane, ABCD1 plays a role in the translocation of very long chain fatty acids (VLCFA) for their damage by β-oxidation. Here, we present five Cryo-Electron microscopy structures of ABCD1 in four conformational states. Combined with functional analysis, we found that substrate and ATP trigger the closing of two nucleotide binding domains (NBDs) over a distance of 40 Å and the rearrangement of the transmembrane domains. Each of the three inward-facing structure of ABCD1 has a vestibule opens to cytosol with variable size. Furthermore, the structure of ABCD1 in the outward-facing state supports that ATP molecules pull the two NBDs together and open the transmembrane domain to the peroxisomal lumen for substrate release. The five structures provide a snapshot of substrate transporting cycle and mechanistic implications for disease-causing mutations.

## Introduction

The peroxisomal half-transporter ABCD1, known as adrenoleukodystrophy protein (ALDP), is the most important member in the ATP-binding cassette (ABC) transporter D subfamily (Kawaguchi and Morita, 2016), and acts as a central role in metabolism of very long chain fatty acids (VLCFAs) (Trinh The et al., 2021). Mutations of ABCD1 commenly leads to an abnormal accumulation of VLCFAs, resulting in X-linked adrenoleukodystrophy (X-ALD), a progressive neurological disorder with variable clinical outcomes ranging from adrenal insufficiency to rapidly progressive and fatal cerebral demyelination (Turk et al., 2020).

Currently, it has been well known that the gene for X-ALD, encodes for a half-ABC transporter, and shared over 900 variants derived from ABCD1 gene mutant, of those, 65% of them affects the protein folding and stability, which results in a marked reduction of ALDP from the membrane. These will cause variable clinic symptom including adrenoleuko-dysfunction, adrenomyelo-neurotrophy and cerebral adrenoleuko-dystrophy (Engelen et al., 2021; Loes et al., 1994; Mohn et al., 2021). However, there is no specific treatment for X-ALD up to now, it is therefore urgent for us to develop small molecules to correct the folding of certain ALDP mutant so as to find effective strategy for the treatment of X-ALD in preclinical trial.

Although the mutant of ABCD1 contributes the accumulation of long-chain fatty acids (VLCFAs) caused by damaged peroxisome β-oxidation to affect the stability of adrenal gland, testis and myelin (Wang et al., 2011), the full information on the ABCD1 structure is lack from enough experimental evidences also. Previously, given knowledge showed that the substrate of ABCD1, as straight-chain saturated fatty acids (Wiesinger et al., 2013), was confirmed at transfection of human ABCD1 reverse transcription DNA into X-ALD skin fibroblasts to restore the oxidative activity of VLCFA β and the VLCFA content in fibroblasts to normal (Braiterman et al., 1998; Flavigny et al., 1999), it was therefore designated as the transporter of VLCFA through the membrane of peroxisome and dependent on the β-oxidation pathway (Kawaguchi et al., 2021). Meanwhile, Van Roermund et al proved that ABCD1 is involved in the transport of VLCFA-CoA through peroxisome membrane to express human ABCD1 in *Saccharomyces cerevisiae* (van Roermund et al., 2008; van Roermund et al., 2011).

Anyway, the bulk of evidences have showed the important role of ABCD1 under both physiological and pathological condition, especially in the metabolism of VLCFA. However, the lack of structural information on the regulation of ABCD1 by binding of substrate and ATP hinders our understanding on the comprehensive transporting process of this undoubtedly important peroxisomal molecule, even under disease condition. Whereas, to analyze the molecular structure of ABCD1 may find the adopted binding site to invent small molecule drug that could be available to the treatment of involved diseases.

Recently, the atomic structure of ABCD4 was determined by single particle Cryo-EM (Xu et al., 2019), revealing the overall architecture of the ABCD family, which provide a novel idea to screen other member of ABCD family. However, ABCD1 shares low sequence identity with ABCD4 (Figure S1), and exhibits different biochemical properties, thereby it may play a solely vital role in organism both physiological and pathological condition, which definitely demand us to know the concrete information from structure biology angle. Here, we present the ABCD1 full structure and exhibit novel data from functional analysis in human beings.

## RESULTS

### Functional characterization of human ABCD1

ABCD1 is a half ATP-binding cassette transporter, and two subunits dimerize to form a functional full transporter in the peroxisome. ABCD1 was overexpressed in HEK293GnTl^-^ cells and purified in detergents for further functional characterization.

We used an NADH coupled ATPase assay for measuring the K_M_ for ATP and the maximal basal turnover rate for the full length ABCD1, the ABCD1-E630Q mutant and the N-terminal truncation mutant which devoids of the N-terminal 54 residues in a similar way to other human ABC transporters (Figure 1). The ABCD1-E630Q mutant exhibits only one thirds of the ATPase activity of the wild type protein, whereas the N terminal deletion mutant (ΔN1-54) have has a higher ATPase activity (Figure 1A,1B and 1E). Thus we choose ΔN1-54 ABCD1 for measuring the ATPase activity stimulated by substrate. Both K_M_ values for ATP (585 ± 55 µM) and the maximal basal turnover rate (29 ± 1 nmol/mg/min) of ΔN1-54 ABCD1 is similar to those reported for ABCD1 in liposome (Okamoto et al., 2018).

**Figure 1.**
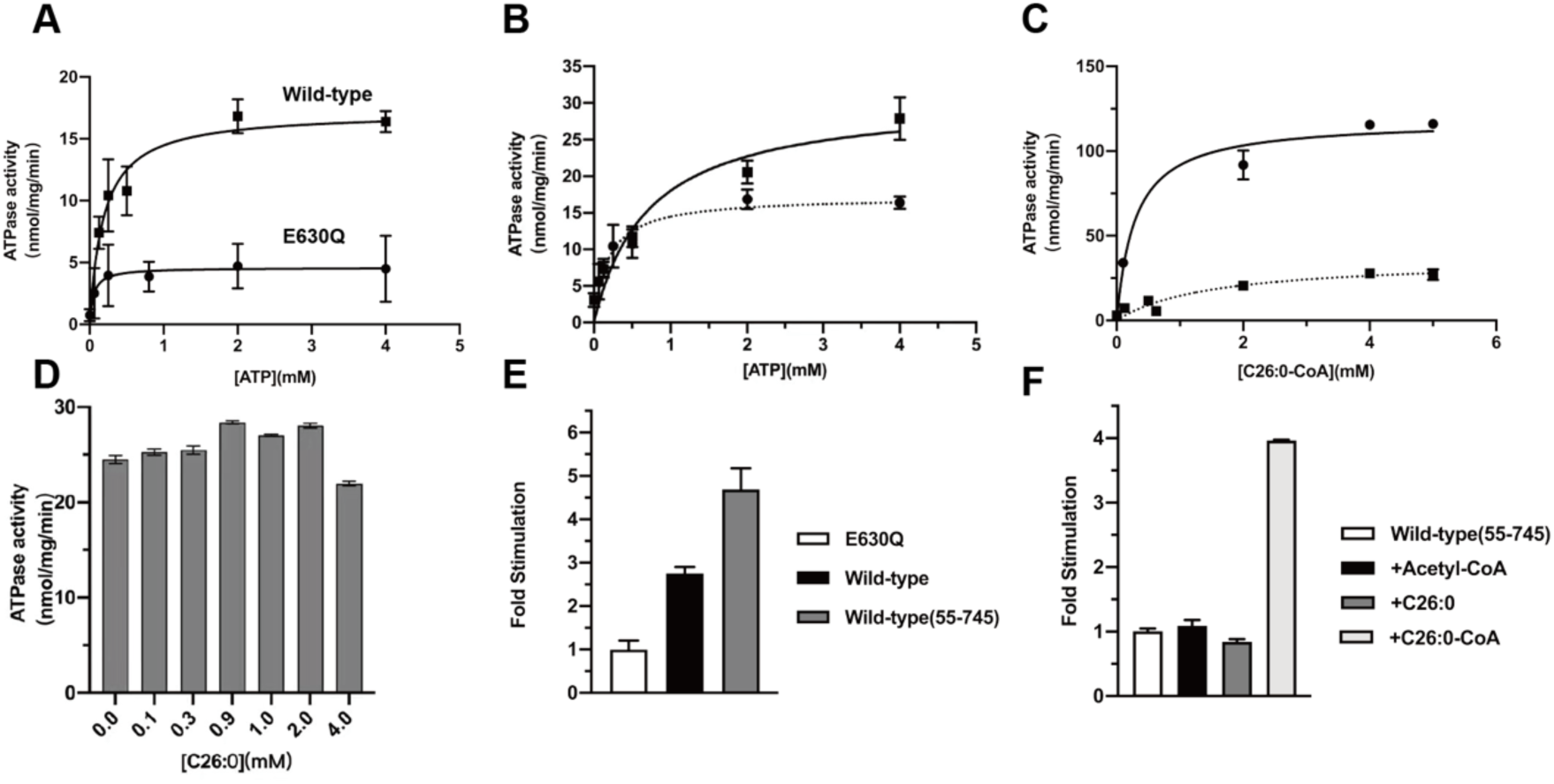
The ATPase activity of human ABCD1 stimulated by substrates. (A) ATPase activity of the full-length wild type and E630Q mutant protein purified in GDN. The full-length wild type shows a K_M_ of 279 ± 18 µM for ATP and a maximal hydrolysis activity was 18 nmol/mg/min. (B) ATPase activity of the truncation mutant devoid of the N-terminal 54 residues. The mutant has a K_M_ of 586 ± 55 µM and the maximal ATPase activity was 30 ±1 nmol/mg/min. (C) ATPase activity of the truncation mutant devoid of the N-terminal 54 residues as a function of C26:0-CoA in the presence of 4 mM ATP. The K_M_ value determined for C26:0-CoA was 286 ± 41 µM. (D) ATPase activity of the truncation muatant devoid of the N-terminal 54 residues as a function of different concentrations C26:0. (E) Normalized ATPase activity of E630Q mutant protein. The stimulation activity of the full-length wild type and the truncation muatant devoid of the N-terminal 54 residues. Maximal hydrolysis activity was selected to calculate the stimulate fold. (F) Stimulated ATPase activity of the truncation mutant devoid of the N-terminal 54 residues in the presence of C26:0, C26:0-CoA and Acetyl-CoA. We selected maximal hydrolysis activity to calculate the stimulate fold. The reported errors represent SD.

We observed significant increase in ATPase activity in the presence of C26:0-CoA substrate (Figure 1C), but addition of either C26:0 or Acetyl-CoA has no stimulation effect (Figure 1D and 1F). The apparent affinity of ABCD1 for its substrate C26:0-CoA is close to that reported for the ABCD1 measured in vesicles, and potting the ATPase activity as a function of C26:0-CoA concentrations confirmed that C26:0-CoA stimulates ATP hydrolysis by 3 fold at saturation concentrations. Moreover, the K_M_ for C26:0 is 0.5mM similar to that reported for plant CTS (Carrier et al., 2019), a plant VLCFA transporter homologue from Arabidopsis thaliana. These validate our experimental system and the detergent purified ABCD1 is reliable and it could be used for later structural analysis.

### Structure overview

To elucidate the mechanism of conformational changes during the transport cycle, we collected datasets on the full length wild-type ABCD1 in the presence of hexacosanoic acid (C26:0), and the ABCD1-E630Q mutant in the presence of ATP and the ABCD1-E630Q mutant in complex with both C26:0-CoA and ATP were performed. Consequently, Cryo-EM reconstructions based on these datasets yielded five molecular structures of ABCD1 in four different conformations: a wide open inward-facing form at 3.78 Å and 3.33 Å for the ABCD1-C26:0 complex as well as ABCD1-E630Q-C26:0-CoA complex (Figure S2 and 4C); an inward-facing intermediate form at 3.34 Å (Figure S4D); and an nearly closed inward facing ABCD1-E630Q, as well as an outward-facing ABCD1-E630Q at 3.3 Å and 2.96 Å in the presence of ATP, respectively (Figure S3D and S3C).

The electron microscopy density map for the wide open inward-facing structure (state 1, Figure 6) is sufficient for us to build all the transmembrane helices and model the NBD regions (Figure S5). The region corresponding to the C-terminal coiled-coil helix has a clear electron density, enabling us to model their side chains. We built 3D structures consisting of residues 64-724 for the ABCD1 in complex with C26:0, and the ABCD1-E630Q in the presence of C26:0-CoA and ATP. Therefore we will use the wide open inward-facing structure to describe the molecular structural features.

The intermediate inward-facing ABCD1 in state 2 has an electron density map with similar quality to that of ABCD1 in state 1, however, the density for the C-terminal coiled-coil is missing. Compared to inward-facing structures (state 1-3), the outward-facing ATP-bound ABCD1 present much better densities corresponding to the nucleotide-binding domain(NBD) region and the cytoplasmic helices of the transmembrane domain (TMD). Thus, we build an atomic model of the NBD and the TMD helices except the loops on the peroxisomal lumen connecting the TMD helices because of their large flexibility and poor electron density maps. In all the above reconstructions, the final structures were refined against the EM maps to get good statistics and reliable stereochemical parameter.

Each human ABCD1 subunit contains one 6-transmembrane helical domain followed by one nucleotide binding domain (Figure 2E and 2F) and two subunits assemble into an isosceles triangle structure in the wide open inward facing state (Figure 2B). One of the two long side from the isosceles triangle was TM bundle 1 that was formed by assembling transmembrane helical structure TM1, TM2, TM3, and TM6 in subunit A with TM4 and TM5 of subunit B. Another long side, also the other TM bundle, is comprised of the TM4 and TM5 of subunit A and TM1, TM2, TM3, and TM6 of subunit B. The two TM bundle, of which angle is the biggest in all ABC family proteins’ structures, formed a large vestibule open to cytoplasm. The vestibule possess high affinity binding site for substrate, which is an important characteristic of ABC exporters. Furthermore, IH2 (Figure S1) between TM4 and TM5 shares same subunit interaction with the NBD of the other subunits *via* Wan der Waals interactions. Whereas, mutations of amino acids on those surfaces alter the normal function of ABC transporters, of these, 80% cystic fibrosis is related to the deletion of 508^th^amino acid in CFTR as mutations in this site influences the 3D structural formation of CFTR protein. The corresponding residue Y559 in ABCD1 interacts with Y292 and E302 of the IH2 of the paring subunit, mutation of each of the three residues in human causes X-ALD disease(Scharschmidt, 1979).

**Figure 2.**
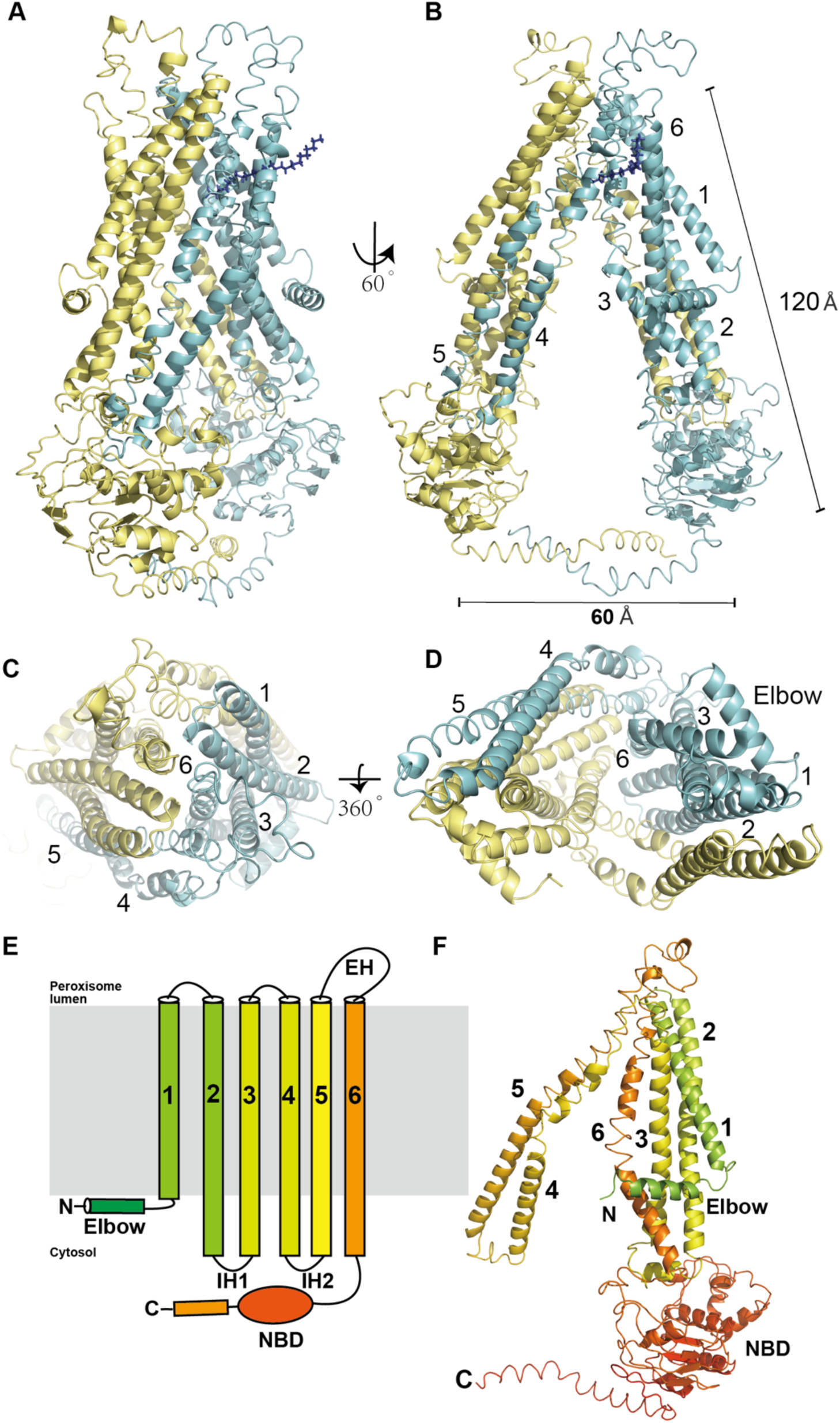
Overall structure of ABCD1 in complex with C26:0 fatty acid. (A) Side view of overall structure of ABCD1 in cartoon representation. The two subunits are colored in cyan and yellow respectively. (B) The transmembrane domain of ABCD1 viewed from peroxisomal lumen and cytosol. (C) Domain structures of ABCD1.

Two NBDs localized in cytoplasm where 30 Å away from cell membrane were separated by a 60 Å long curved C-terminal coiled-coil, therefor formed the two angles of isosceles triangle. Interestingly, what we can imagine is, the two NBDs should cross an amazing 30 Å area before coupling tightly together with the presence of Mg^2+^, and reasonably influences the alteration of the conformation in transmembrane area of ABCD1 from inward-facing one to outward-facing one.

### Substrate recognition

ABCD1 was found both in conformation 1 and 2, whose C-terminal tail formed coiled-coil dimer structure, parallel separating two NBDs. We observed both the C26:0 and C26:0-CoA substrate bind to a similar position located at the interface formed by 4-helix, TM3, TM4, TM5 and TM6 in the TM domain, half away from the cytosol (Figure 3 and S7). The hydrophilic head group protrudes into the vestibule with their tails buried into the fenestration formed by TM3, TM4, TM5 and TM6(Figure S7A and S7B). In the case of C26:0, the carboxylic group locates near the inner surface of the vestibule formed by residues E400, S404, E408, W339 and S340 (Figure 3D), and the tail lies in the fenestration via hydrophobic interactions. The C26:0-CoA substrate binds to the same position of ABCD1-E630Q as C26:0 in the wild type ABCD1 structure. Figure 3H shows the interactions between the head group of CoA and the protein, which involves mainly hydrophobic and hydrogen bond interactions with the surrounding residues. W339 and M335 from TM5 pack with the adenine moiety from one side via hydrophobic interactions, whereas R401 and S404 stabilize the 3’-phosphate groups from the other side via hydrogen bond interactions. The pantetheine arm extends into the fenestration, encircled by hydrophilic and hydrophobic residues such as S226, L230, F252 and L249. The fatty acid tails of C26:0-CoA appears to be flexible as their density is less well defined.

**Figure 3.**
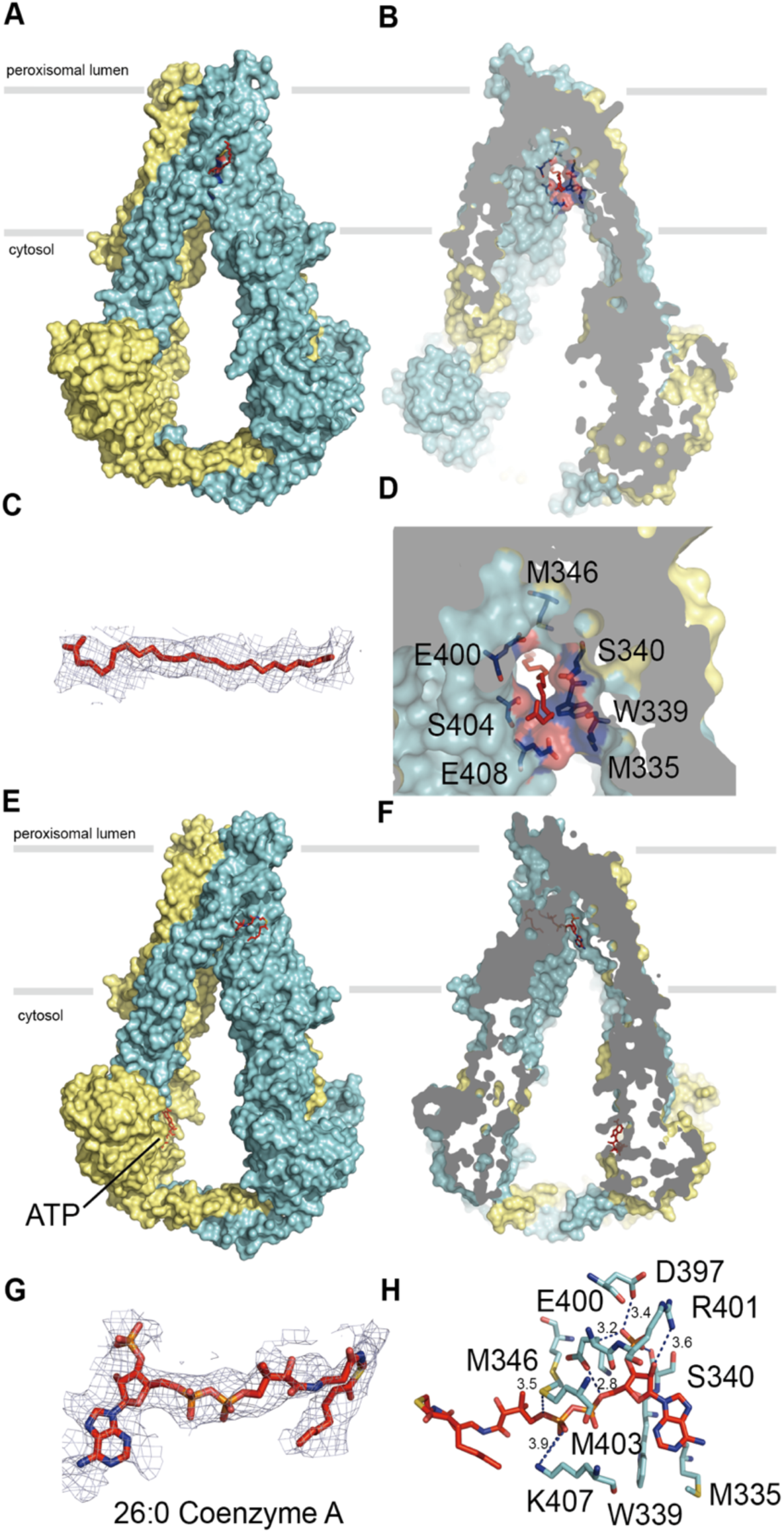
The inward-facing structure of ABCD1 has a cytosolic vestibule that houses the substrate binding site. (A) and (B) Surface representation of ABCD1 shows C26:0 binds to a vestibule that penetrates half a way into the lipid bilayer. (C) and (D) Electron density map of C26:0 substrate and zoomed in view of the substrate in the same orientation as (B). (E) and (F) The overall and slice views of ABCD1-E630Q in complex with C26:0-CoA as surface representation. (G) and (H) EM density of parts of C26:0-CoA and zoomed in view of the CoA binding site.

A unique feature of ABCD1 in state 1 in the present study is that the distance between the two NBDs approached 50-60 Å, which beyond our anticipation before this test. Notably, the distance between ATP combining site (Walk A motif) and signature motif was 38 Å (Figure 6A), which is accordant with previous report, and the distance of the two NBD domains of CFTR under close state also uniquely attained 45 Å (Chaves and Gadsby, 2015). In principle, combined with previous experience and reports, the intracellular level of ATP could not make two NBD molecules separate such far in distance. Whereas, the presence of the coiled-coil helix in the wide open inward facing conformation probably configures the protein ready for binding the long-chain fatty acid substrate.

Among other ABC transporting proteins, such as ABCC1-apo structure, the distance between two NBDs domains of ATP binding sites are so far, as reported previously, attained 35 Å of mean level (Johnson and Chen, 2017). However, under this condition, the two NBDs domains of ATP binding sites were uncommonly located opposite in unparallel style. In this study, the parallel opposite location of the two NBD domains of ATP binding sites may be attributed to the C-tail of ABCD1 that made the two NBD domains of ATP binding site lined up in a relatively parallel location.

Structural comparison of the two NBDs domains in the substrate-bound structure reveals that the very end of the C-terminal coiled-coil from one subunit interacts with the other NBD differently. In one NBD domain, E722 from the coil helix is positioned closely to P508 from the Walker A motif (Figure S5B and S5C), whereas E722 in the other subunit moved away from residue P508. Astonishingly the sequence of the Walker A motif (GPNGCGKSS) (Figure S1 and S6G) is conserved among the ABCD1, ABCD2, and ABCD3, but different from that of ABCD4. Furthermore, we are definitely to define an ATP molecule binds to the Walker A motif of the same NBD domain in the C26:0-CoA bound structure (Figure S6B), of which, the E722 makes close contacts with P508. We suspect that binding of substrate induces asymmetrical conformational changes of the NBD, which propagates to the C terminal coiled-coil.

### The inward-facing conformational intermediate of ABCD1

The pro of ABC family is generally considered that when binding to the TM domain of substrates, it is vulnerable for combining its prime protein state with ATP molecules and hydrolyzing them. In this study, we also conducted the experiments to elucidate the role of C26:0 and C26:0-CoA in promoting the ABCD1 ATPase. C26:0-CoA but not C26:0 significantly elevated the vitality of ABCD1 ATPase activity by about 3 fold (Figure 1C). Furthermore, we performed the test referred to the other methods in which the ABCD1-E630Q mutation, ABCD1 inhibition and hydrolysis of ATPase was set up respectively in the typical biochemistry experiments, the ABCD1-E630Q mutant reduces the ATPase activity to one half of the wild type ABCD1. Therefore, based on these, we attempted to investigate the structural changes under the condition of ABCD1 with the addition of substrate compounds.

Three dimensional reconstructions of the complex of ABCD1-E630Q mutant with C26:0-CoA and ATP yielded an additional inward-facing intermediate conformation(Figure 6C and 4), in which, the density of the C-terminal coiled-coil is definitely missing. Under the special state, TM4 moves towards TM3 and TM6, closing the fenestration that extends from the TM4-TM3 interface to the substrate binding site near W339 residue. Consequently, the side chains of W339 from TM4 and R401, K407 from TM6 close the open court by forming cation-π interactions, without the possibility to combine with the CoA part into the position near W339 (Figure 4B). The undergoing movement of the TM bundles forms a smaller angle between the two TM domains with a slit surrounded by residues N256, R259, T416, R411 and F252 (Figure 4C). Its position is closed to the N-terminal region of the elbow, which lies on the outside of TM6 and TM3 helices. However, we could not define the electron density for the N terminal 54 residues preceding the elbow. Previous studies suggest that mutation of residues in the elbow affect the folding of ABCD1, resulting in lower levels of protein in body tissue. The tight interaction between the elbow and TM6 could make the elbow remain attached on the TM6 during the movement of the inner parts of the TM domains during transport cycle. Therefore, the preceding the N-terminal 54 residues region would have a higher probability to affect the opening of the slit or perturb the exposing of the substrate-binding site. In this kind of conformation, two NBD domains moved closer to each other, resulting in the disruption of the dimerization of coiled-coil in C-tail. We also identified one ATP molecule binds to the Walker A motif in the NBD domain of one subunit (Figure S6C). Moreover, deletion of the N-terminal 54 amino acids significantly increase the ATPase activity of ABCD1 with and without the addition of C26:0-CoA compared to the full-length protein (Figure 1B).These results suggest that the N-terminal region preceding the elbow has an inhibitory role of ABCD1, probably regulates the opening of the two bundles for exposing the substrate-binding site.

**Figure 4.**
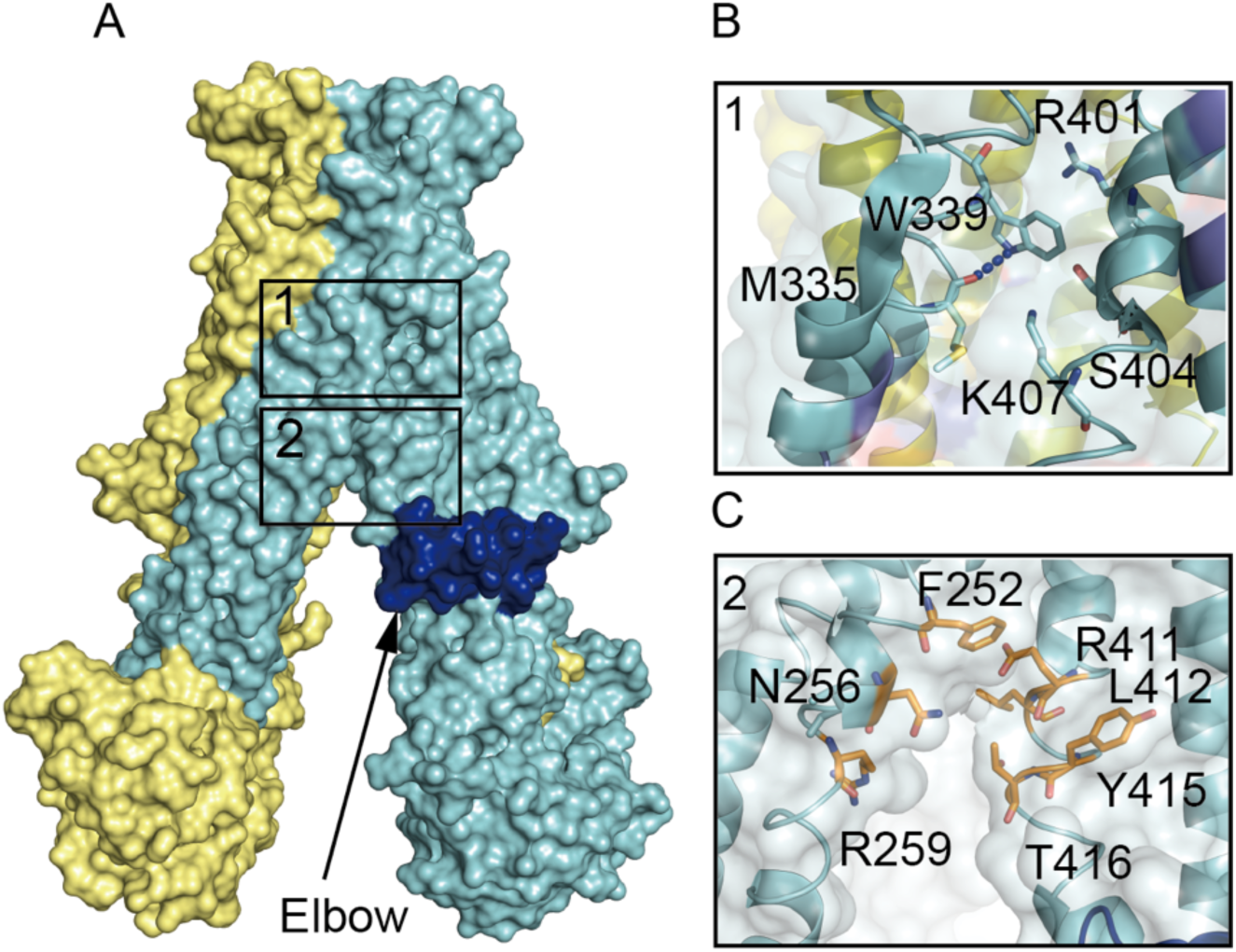
The inward-facing intermediate structure of ABCD1. (A) Surface representation of the inward-facing intermediate structure of ABCD1. (B) The substrate binding site near W339 closes. (C) Potential modulation site near the elbow.

### ATP binding drives the inward-to-outward conformational transitions

It has been found that ABCD1 is a kind of half transporter, needed for the dimerization of two molecules to form functional molecular. Above, we described that C-tail could form dimers, and thereby make the binding sites of two ATP molecules align in parallel style. With the help of Cryo-electron microscopy, we confirmed that ABCD1-E630Q molecules could combine with ATP, in K_M_ of 0.4 mM, therefore reported a novel kind of inward facing conformation that could bind to two ATP molecules, displaying as “hand in hand” confirmation for two NBD domains linked by N509 residues from both subunits (Figure 5H and S6D), but the two NBD domains twisted, with complete separation of Walk motif and signature motif of both the ATP binding sites. But ATP molecules only bind to the Walker A motif of either NBD domain and the Mg^2+^ ion could not be defined from the electron density map (Figure S6D). Without the interaction with the signature motif and the absence of Mg^2+^ ion, the ATP molecules could not be hydrolyzed normally. Structural comparison of the ATP-binding sites in ABCD1 between state 2 and 3 shows that N509 switch from the interaction with H659 in state 2 to N509 from the paired subunit in state 3 (Figure S6D). The mutation of N509 residues of the Walker A motif can leads to diseases. While in ABCC1 and ABCD4, the corresponding residue change to Valine and Threonine respectively (Figure S6G), ie. different from those of ABCD1, so it is needed for both the Walker A motif and signature motif approaching to form necessary ATP binding sites, it may be the possible explanation of the phenomenon in our study that under the condition of combination with C26:0, and the distance of two NBD domains was 40 Å, it was feasible for them to bind with ATP molecules. These ATP binding sites of unique reflects the distinct molecular mechanism with which ABCD1 plays its roles. The fact that ATP molecules bind to three states of ABCD1 without hydrolysis suggests it modulates the activity of the protein by regulating key residues such as H659, N509, and P508.

**Figure 5.**
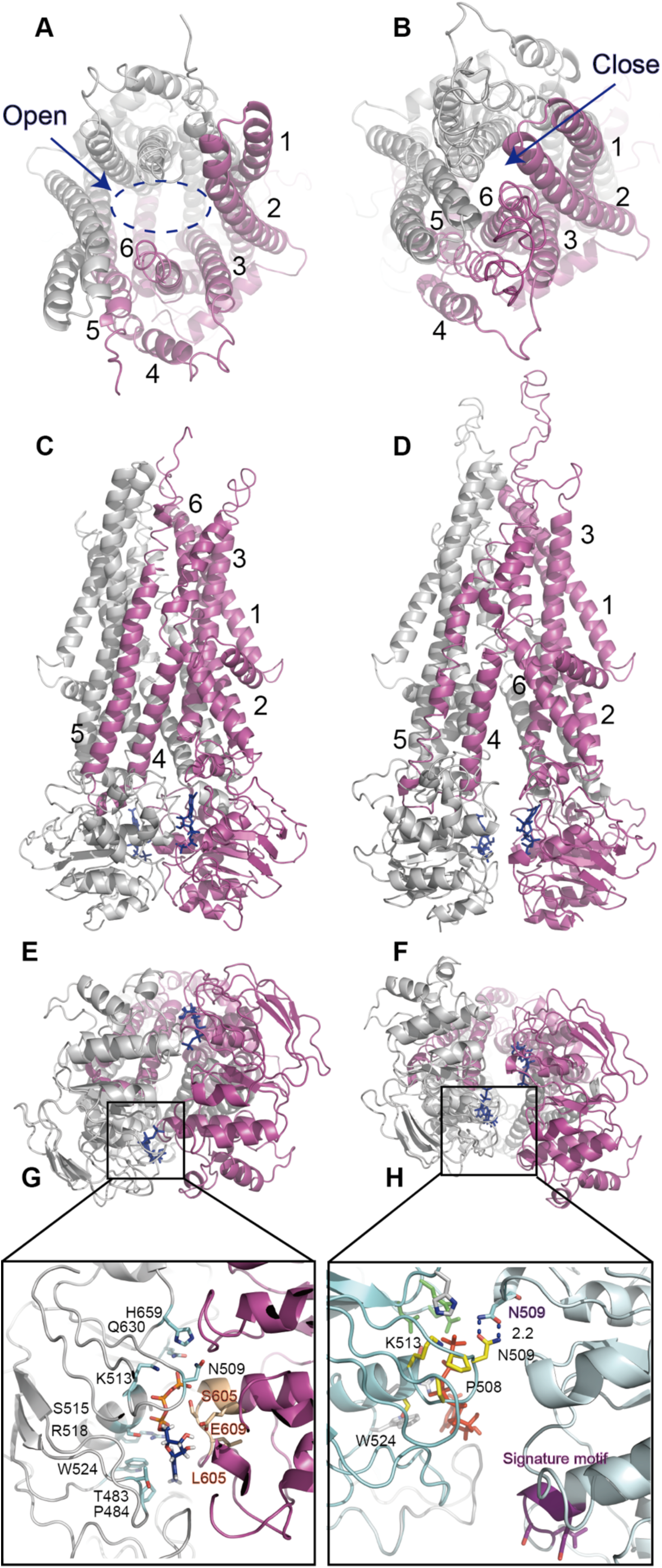
The ATP-bound inward-facing and outward-facing structure of ABCD1. (A), (C), (E) and (D) Represent the cartoon structure of ABCD1 in complex with ATP in the outward-facing state. (A) and (E) Viewed from the peroxisome lumen side and the cytosolic, while (C) shows the side view. (G) Zoomed in view of the ATP binding site. (B), (D), (F), and (H) Capture the cartoon representation of ABCD1 in complex of ATP in the inward-facing state viewed from a similar orientation as the outward-facing structure on the left.

However, in the ATP-bound outward state, the “hand-in-hand” interaction between both N509 is disrupted and each N509 form two hydrogen interactions with the carbonyl of S633 and main chain nitrogen of V635, to stable the canonical ATP-binding site (Figure S6E). These hydrogen interactions were found in our observation and it is probably important to maintain the normal function of ABCD1 as mutation of three residues can lead to disease. The adenine moiety of ATP molecule is sandwiched between P484 and W524 of one NBD and V604 from the other NBD. The phosphate groups of ATP form electrostatic interactions with the Walker motif, the Q-loop of one NBD and the signature motif of the other NBD. ATP molecule acts like a glue to staple the two NBDs together, thereby bring two bundles of TMDs together and forms new interactions between one NBD and the IH1 and IH2 of the paring TM bundle (Figure S6F). Therefore, these interactions likely couple the conformational changes induced by binding of ATP to the transmembrane domain and drives the transition of the inward-facing to the outward-facing state.

We can extrapolate the conformational changes of ABCD1 during substrate release by comparing ATP-bound inward-facing and outward-facing structures. The closure of NBDs by binding of ATP bring the two halves together as indicted by shortening of the distance between the cytoplasmic parts of TM3 and TM4 from 16Å in the inward-facing state 3 to 7Å in the outward conformation (Figure 6).The cytoplasmic part of TM5, directly linked to TM4 through the inner IH2 and tight hydrophobic contacts with TM3, moves towards to the opposite halve and the close the deep vestibule that opens to the cytosol in the inward-facing conformation. The inner helix of TM2, which connects to TM3 through IH1 and tight contact with TM3, also moves along with TM3 to help seal the vestibule. The elbow embraces the surface of TM 3 and TM6 like a string though tight hydrophobic interactions and may play roles in regulating the conformational change of TM 3 and TM6. The extracellular ends of the TM helices rotate outwardly and open the transmembrane vestibule to the peroxisomal lumen. The outward-facing structure of ABCD1 has a remarkable big vestibule which seals at the deep cytoplasmic side with electrostatic interactions between charged residues (Figure S7G). These residues at the bottom of the vestibule include R276 and K276 from TM6, Q195 and E195 from TM2, which is underneath the elbow (Figure S7G). The inner surface of the vestibule is decorated with a number of charged residues, which may exclude the binding of the hydrophobic tails of the long chain lipid and help release it into peroxisome. Like the outward-facing structure of ABCC1, we could not identify any substrate in the vestibule, which may not possess a binding site with affinity high enough for us to capture the substrate. This outward-facing structure provides a glimpse of the substrate-release state of ABCD1.

**Figure 6.**
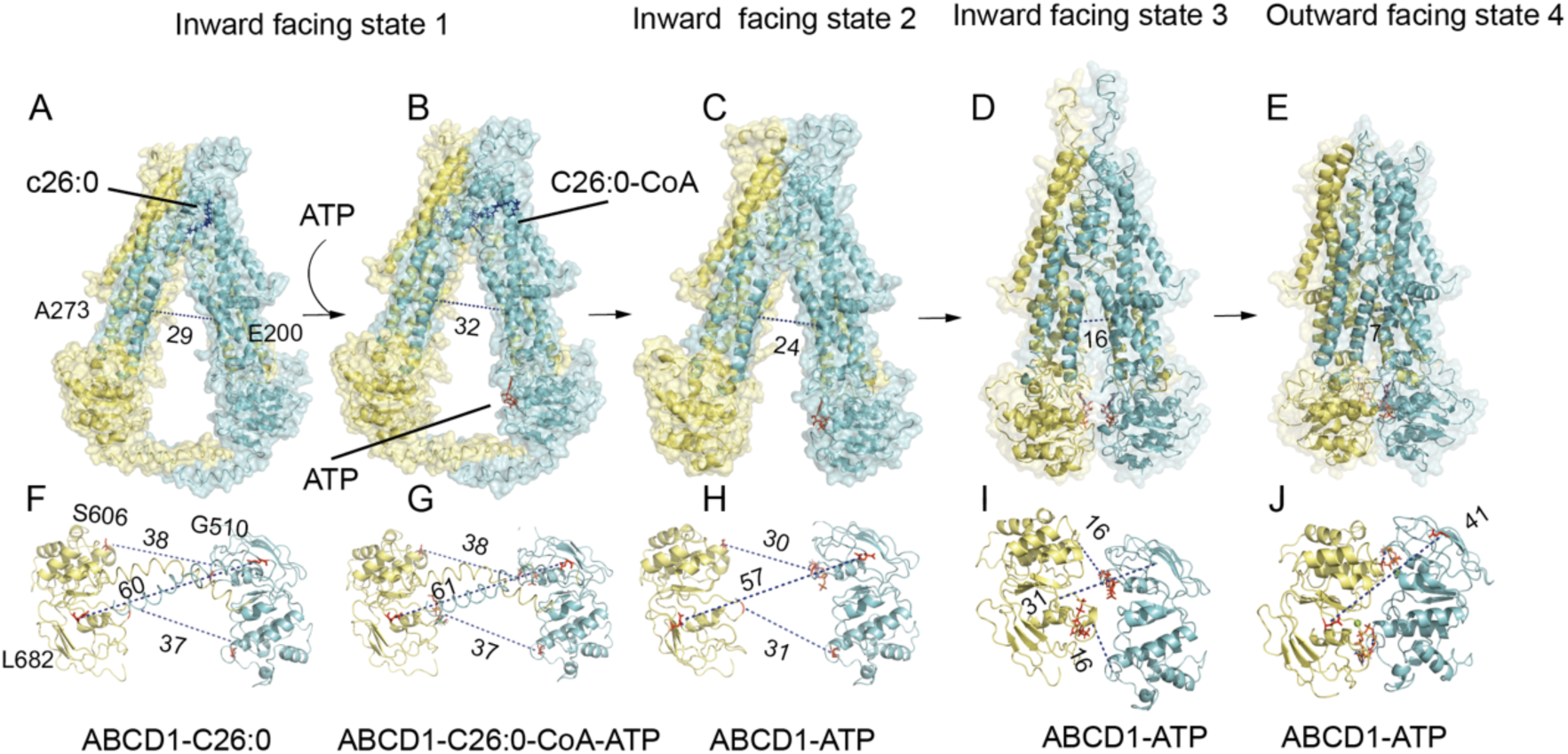
The three inward-facing and one outward-facing states of ABCD1. (A) and (B) Surface representation of ABCD1 in the inward-facing state 1, characterized by the presence of C-terminal coil. (C) The structure of ABCD1 in the inward-facing state 2 has a shortening distance of two NBDs compared to that of state 1. (D) ABCD1 has a narrower vestibule opens to the cytosol when it has been captured in the inward-facing state 3. (E) ATP staples the two NBDs and the outer segments of the TMD are pealed open to the peroxisome lumen. (F)-(J) Measuring the distance between the two NBDs according to different states of ABCD1 on the above.

## Discussion

### Conformational changes during the translocation process of ABCD1

In the present study, we captured three inward-facing conformations and one outward-facing conformation (Figure 6). We measured the distance between S606 of the signature motif of one NBD and G510 of the Walker A motif of the other NBD in all the above states except for the outward-facing conformation, in which both residue comes together to bind the ATP molecule. E682 at the end of the final β-strand of a typical NBD is another residue we choose for measuring the relative position of the two NBDs in different states. To analyze the conformational changes in the transmembrane domain, we measured the distance between D200 of TM3 and A273 of TM4, both of which move close to a distance of 7 Å in the outward-facing state upon binding of ATP.

Comparing these structures, we found that the two ATP-binding sites were 38 Å apart, TM3 and TM4 were 30Å apart in state 1 (Figure 6F). In this state, the C-terminal of the coiled-coil of one NBD closely attaches to the Walker A motif of the other NBD, thus N509 at the Walker A motif can stabilize the phosphate group of the ATP molecule. Upon binding of substrate and ATP, the two NBDs moves towards each other along the axis of the C-terminal coiled-coil, and thus the distances between the two separated ATP binding sites are shortened by 7 Å in state 2 (Figure 6C). The C-terminal coiled-coil is missing in this state, otherwise it would block the movement of the two NBDs. Without the interaction with the C-terminal coiled-coil, N509 of the Walker A motif interact with both H659 and the phosphate group of ATP (Figure S6B). Furthermore, the two bundles of transmembrane domains moves closer too, with a shortening distance of about 6 Å (Figure 6C). In state 3, the two NBDs appear to be very close with forming the new interaction of N509 from both subunits (Figure 6I and Figure 5H). However, the ATP binding sites are completely separated, with a distance of 16Å between S606 from one NBD and N509 from the other NBD (Figure 6I). In state 3, ATP binds to the Walker A motif with its phosphate group stabilized by K513, Q630, D629 and Q544 (Figure S6D). Although ATP molecule can bind to one NBD independently, it cannot be hydrolyzed without the interaction of the signature motif from the other subunit and Mg^2+^ ion. Therefore, the binding of one ATP molecule to A motif Walker A of one NBD probably plays a role in pulling the signature motif from the other subunit. The distance between TM3 and TM4 are shortened further by about 8 Å compared to their distance in state 2 (Figure 6D). However, the inner halves of the transmembrane segments have not seal the vestibule yet at the cytosolic side in this state. Until the two NBDs staples together (Figure 6E and 6J), the conformational changes occurred at the NBD-TMD and the inner halves of the transmembrane segments propagate to the outer halves of TMD domains and open the vestibule to the peroxisomal lumen, subsequently the substrate release into peroxisome. The hydrolysis of ATP molecules would reset the transport cycle. In conclusion, our structural analysis indicates binding of both substrate and ATP molecule would stimulate transporting activity of ABCD1 more efficiently, because of the two NBDs are separated by a large distance by the C-terminal coiled-coil. Binding of ATP would release the brake acted by the C-terminal coiled-coil by influencing the Walker A motif which coupling to the C-terminal of the coiled-coil of another subunit.

### Structural basis of disease-causing mutations

ABCD1 is the gene with the most mutated loci found in the ABC family and most of these mutations are pathogenic. We analyzed the database (https://adrenoleukodystrophy.info/) or previous literature for mapping of these mutations, where the red-labeled mutation cause protein misfolding and degradation, and found that most of these mutation sites are evenly distribute along the full-length protein (Figure 7). A total of 1226 mutations have been reported with 826 mutations cause disease. Among these mutations, most of them influence the folding and stability of the protein, resulting in reduced levels of protein in patient, with only 30 mutations of non-functional protein, although they reach to a normal level in human tissues. Considering a large conformational changes during transporting process, we suspect that the mutation sites would prefer to affect one of the conformational states (inward-facing vs outward-facing), and therefore we map their positions separately onto both the inward-facing and outward-facing structures according to their roles in maintaining one of the states. For example, the inner helix of the TM3 and TM4 interact only in the outward-facing state, therefore we map the mutations on this region in the outward-facing conformation. Of course there are some sites of residues which interact all the time during the transporting cycle. We found that most of the interacted-residues located at key positions of ABCD1 such as the TM domain intersection, NBD-TM interfacing, ATP-binding site, and substrate binding site, and mutation of these positions are associated with disease, and we labeled them as red sphere in the inward-facing conformation or the purple sphere in the outward-facing conformation. In contrast, there were mutations that did not affect protein expression levels, but these proteins were loss of function, and we have labelled these mutations blue and orange separately on the inward-facing and outward facing structures (Figure 7A and 7B), which can develop activity-regulating molecules against these mutations for clinical in the future.

**Figure 7.**
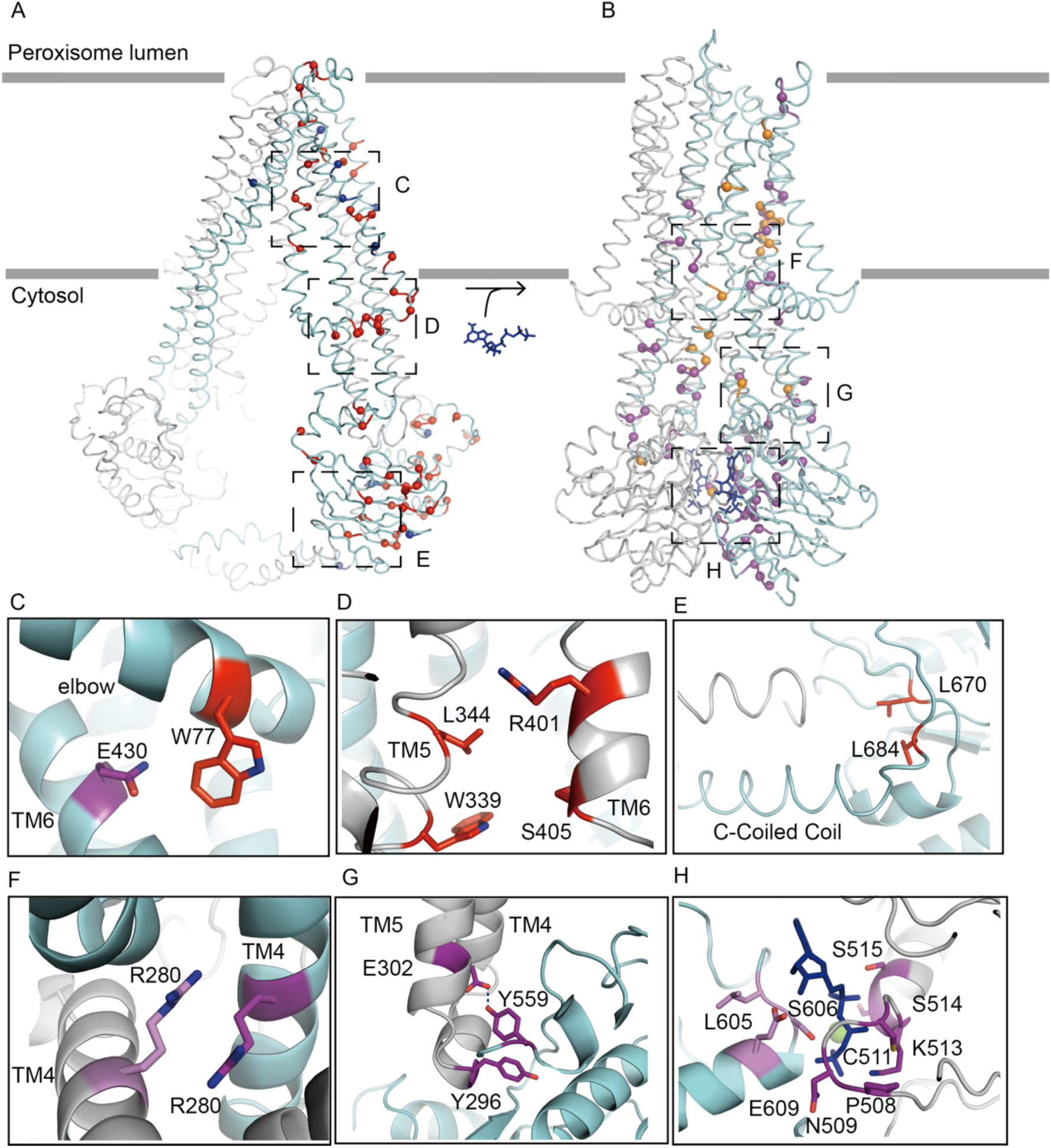
Mapping the locations of disease causing mutations using the inward-facing and outward-facing structure of ABCD1. (A) Red balls represent the deleterious mutation causing the reduced expression of ABCD1 in human tissues, which were chosen because they mostly form interactions in the inward-facing structure. Blue balls mean their mutation does not affect the level of protein in human tissues but results in normal expressions of protein with functional defects. (B) Location of mutations on the structure of ABCD1 in the outward facing state. Purple balls represent the deleterious mutants and orange balls shows mutation that affect the function of ABCD1 but does not cause a folding problem. (C-H) Roles of representative residues by three-dimensional structural analysis.

The number of mutations is incredibly bigger than its total number of amino acid residues. Among an amount of 826 missen mutations, only 218 mutations have been studied to affect the folding or function of ABCD1 and we mapped them separately on the inward-facing and outward-facing structure. Therefore understanding the action of each residues and cluster of residues in maintaining a certain conformation or triggering alteration of conformational states are valuable in developing correcting molecules and potentiators for the treatment of X-ALD. The FDA approved drug ivacaftor for the treatment of cystic fibrosis increases the open probability of wild type CFTRs and mutant CFTR (Liu et al., 2019; Zhang and Chen, 2016). A similar drug development strategy may be possible to develop activators for mutant ABCD1, as a number of 30 mutant influences the activity of ABCD1.

Structural analysis of the inward-facing and outward-facing conformations of ABCD1 reveals that a number of clusters of interacting residues, mutation of which can cause disease. Figure 7C-E presents the examples of interacting residues on the inward-facing state of ABCD1. W77 on the elbow interacts with E430 from TM6 (Figure 7C), mutation of either of them result in low level expression of ABCD1 (Matteson et al., 2021; Park et al., 2014). The fact that the N-terminal deletion mutant enhance ATPase activity of ABCD1 indicates W77 may have role in regulating the activity of ABCD1 via direct contact with TM6. On the other hand, L684 positioned at the very beginning of the C-terminal coiled-coil (Figure 7E), its mutation to proline may affect the correct orientation of the coiled-coil helix, decoupling its interaction with the Walker A motif of the NBDs. Furthermore, mutations to a variety of residues have been reported on positions of W339 and R401(Chu et al., 2015; Coll et al., 2005; Kumar et al., 2011; Wichers et al., 1999; Zhang et al., 2019), which situates at the substrate binding site (Figure 7D). In addition, a large number of interaction appears at the outward facing conformation of ABCD1. The inner helix of TM4 on separated TM bundles only moves close to each other in the outward-facing conformation and R280 on both helix forms salt bridge interaction at the very bottom of the outward-opening vestibule. Mutation of R280 to Cystine may lock the protein in an outward-facing conformation (Coll et al., 2005; Lan et al., 2011), further structural studies of R280C will be needed to clarify its influence on the function of the protein. A large number of cases have been reported at the S606 and G512 position from signature motif and the Walker A motif respectively (Gartner et al., 2002; Kumar et al., 2011; Zhang et al., 2011), which interacts with the ATP molecule at the outward facing conformation. A large number of mutations occurred at the ATP binding site, which may affect the activity of ABCD1. Lastly the mutation of Y559(Karkar et al., 2015), a cystic fibrosis ΔF508 mimic(Korenke et al., 1997), may affect the interaction of the NBD of one subunit with the IH2 of the other one.

In summary, we present a series of conformations of ABCD1 from the wide open inward-facing vulnerable for binding of substrate to the outward-facing state for releasing the substrate. Both the substrate and ATP stimulate the activity of ABCD1 through the TMD and NBD domains. A C-terminal coiled-coil, a unique feature of ABCD1, regulates the activity of the protein by direct contact with the Walker A motif of the other NBD. Furthermore, we develop a map of a large number of mutations on the inward-facing and outward-facing structures. These structures not only provide a snapshot of conformational changes during the transport cycle, but also offers a basis to understand human X-ALD and facilitate drug development.

## Acknowledgments

We would like to thank the Kobilka Cryo-Electron Microscopy Center, The Chinese University of Hong Kong, Shenzhen for our Cryo-electron microscopy and we would be grateful to Zheng Liu for his help of making EM samples, Cryo-EM grids, taking and analyzing EM images, operate cryo-electron microscopeand the National Facility for Protein Science in Shanghai (NFPS). This work was supported by the National Natural Science Foundation of China Grants 31770897, 81801294; National Key Research and Development Project 2017YFA0504300; the International Science and Technology Innovation Cooperation Project of Sichuan (No.2021YFH0141).

## Author Contributions

C. X. carried out constructs and purified proteins. C.X. and MH.SH. measured the ATPase activity. L. T. Z.L, and LN. J. collected, processed, and interpretated cryo-EM data. L.T. and Z.L, LN.J conceived, designed, and supervised the study and wrote the manuscript.

## Data availability

The Cryo-EM structures reported in this paper are PDB: ####, ####, ####, #### and ####.

### Methods Cell culture

HEK293S GnTl^-^ suspension cells were cultured in Freestyle 293 medium (GIBCO) at 37°C, supplemented with 8% CO_2_, 2% FBS and 80% humidity.

### Expression and purification of human ABCD1

Full-length Human ABCD1 gene was synthesized by General Biosystems company. The Human ABCD1-E630Q was introduced using a standard two-step PCR. Both wild type and mutant gene were cloned into a vector with an N-terminal Flag tag (DYKDDDDK). HEK293S GnTl^-^ cells were transiently with the expression plasmid and polyethylenimines (PEIs) (Polysciences) when cells density reached 3×10^6^ cells per mL. For 3 L HEK293S GnTl^-^ cell culture, 5g plasmids were premixed with 10 g PEIs in 300 mL fresh medium for 5 min, then the mixture was added to 3 L cell culture. The transfected cells were incubated at 37°C for 72 hr and harvested.

For purification, after centrifugation at 4000 rpm for 30 min, the cell pellets were resuspended and solubilized for 3 hr 4°C in the lysis buffer containing 150 mM NaCl, 25 mM Tris pH 8.0, 2 mM MgCl_2_, 20% (v/v) glycerol, 1% (w/v) n-Dodecyl-β-D-Maltopyranoside (DDM), and 0.1% (w/v) cholesteryl hemisuccinte (CHS) supplemented with protease inhibitors (1 µg/mL leupeprin, 1 mM benzamidine, 100 µg/mL soy trypsin inhibitor, 1 mg/mL pepstatin, 1 µg/mL aprotinin and 1mM PMSF). After centrifugation at 40000 rpm for 1hr, the supernatant was applied to anti-FLAG M2 affinity gel (Sigma Aldrich) at 4°C for 2 hr. The resin was packed into a column and washed with wash buffer contanining 150 mM NaCl, 25 mM Tris pH 8.0, 2 mM MgCl_2_, 5% (v/v) glycerol, 0.02%(w/v) DDM (Anatrace), 0.002% cholesteryl hemisuccinte (CHS, Anatrace).

The protein was eluted with the wash buffer supplemented with 0.2 mg/mL FLAG peptide, the flow through was collected and concentrated by a 100-kDa Centricon (Milipore), and further purified by Superose 6 size-exclusion column (GE Healthcare) equilibrated in 150 mM NaCl, 25 mM Tris pH 8.0, 2 mM MgCl_2_, 0.04% GDN. Peak fractions were pooled and concentrated for ATPase assays or Cryo-EM experiments.

### ATPase activity assay

The ATPase activity of human ABCD1 was determined using an enzyme-coupled reactions that aoxidize NADH after ATP hydrolysis to ADP(Scharschmidt, 1979). The reaction buffer contains 0.04% GDN,150 mM KCl, 50 mM Hepes pH 7.5, 10 mM MgCl_2_, 2 mM DTT, 60 µg/mL pyruvate kinase, 32 µg/mL lactate dehydrogenase, 4 mM phosphoenolpyruvate, and 150 µM NADH, and a final concentration of 0.8 µM protein was added. Reactions were started by the addition of ATP and incubated at 37°C. The NADH consumption was calculated by detecting the fluorescence at λ_ex_ = 340 nm and λ_em_ = 445 nm using CLARIOstar microplate reader (BMG). Rates of ATP hydrolysis were determined by the consumption of NADH and the fluorescence loss of NADH was calculated by the standards of NADH. V_max_ and K_M_ were calculated using GraphPad Prism by fitting Michaelis-Menten equation. Then, the substrates were added into the reaction mixture. The mixture was pre-incubated 10 min before the addition of 4 mM ATP to initiate the reaction.

### Cryo-EM sample preparation and data collection

Purified ABCD1 were concentrated to 3 mg/ml follows by frozen sample preparation, 3 µL protein sample was applied to glow-discharged holey carbon grids (Quantifoil Au R1.2/1.3, 300 mesh). After incubation on the grids at 4°C under 100% humidity, grids were then plunge-frozen in liquid ethane using a Vitrobot (Thermo Fisher). CryoEM datasets were acquired on a Titan Krios microscope operated at 300 kV with a K3 Summit direct electron detector. Images were recorded with Serial EM (Mastronarde, 2005) with magnification was 105 K and pixel size was 0.83Å Defocus range from -1.5µm to -2.5µm. Each micrograph was dose-fractionated to 50 frames under a dose rate of 1.33 e^-^.

### Cryo-EM data processing

For ABCD1-ATP sample, 4152 micrographs were obtained after careful filtering by patch motion correction and patch CTF (Cryosparc) (Alp Kucukelbir1, 2013) from 4290 movies. 2000 particles were manually picked follows by two runs of 2D sorting for automatic particle selection template. 14993572 particles were selected based on the 2D sorting template and 636284 particles were obtained finally for next-step calculation after several runs of 2D sorting for removing bad particles. Initial models in this study was generated by Cryosparc, of which 2188534 particles were filtered by heterogeneous refinement and Non-Uniform Refinement then two conformations were obtained of which resolution were 3.30 Å and 2.96 Å, respectively (FSC was 0.143).

For ABCD1-C26:0 dataset, 8757 movies were obtained firstly and then 8324 micrographs were selected after careful filtering by patch motion correction and patch CTF (Cryosparc). This was followed by manually picked 2000 particles for 2D classification in twice for generating automatic particle selection template. After removed bad particles after several runs of 2D classification of 4224743 particles, 731411 particles were selected for next calculation. Subsequently, two of the initial models (505844 particles) that generated by Cryosparc were filtered by heterogeneous refinement and Non-Uniform Refinement and then two conformations were obtained of which resolution were 3.78 Å and 4.36 Å, respectively.

For ABCD1-C26:0-CoA-ATP sample, 10159 movies were collected before delivering to patch motion correction and patch CTF estimation (Cryosparc) and then selected 9622 micrographs carefully. Manually picked 2000 particles were performd, followed by two runs of 2D classification for generating automatic particle selection template. Based on the template, 6835392 particles were obtained and filtered by times of 2D classification for removing bad one and then 63010722 particles were selected for next steps. The initial models generated by Cryosparc were filtered by heterogeneous refinement and Non-Uniform Refinement and then 8 conformations were obtained of which resolution were 2.86 Å, 3.12 Å, 3.25 Å, 3.29 Å, 3.41 Å, 3.34 Å, 3.33 Å, and 3.97 Å, respectively.

All resolutions were estimated by applying a soft mask around the protein density and the gold-standard Fourier shell correlation (FSC)=0.413 criterion. ResMap was used to calculate the local resolution map.

### Model building, refinement and validation

De novo atomic model buildings were conducted in Coot (Emsley and Cowtan, 2004), and amino acid assignment was achieved based mainly on the clearly densities for bulky residues. Phenix (Adams et al., 2010) was used to refine the model against the electron density map. Iterative cycles of refinement and muanual rebuilding were carried out with phenix and refmac (Brown et al., 2015) with secondary structure restraints. Models were validated using previously described methods and Figures were prepared using UCSF Chimera (Pettersen et al., 2004) and PyMOL (http://www.pymol.org/).

---

**Figure S1.**
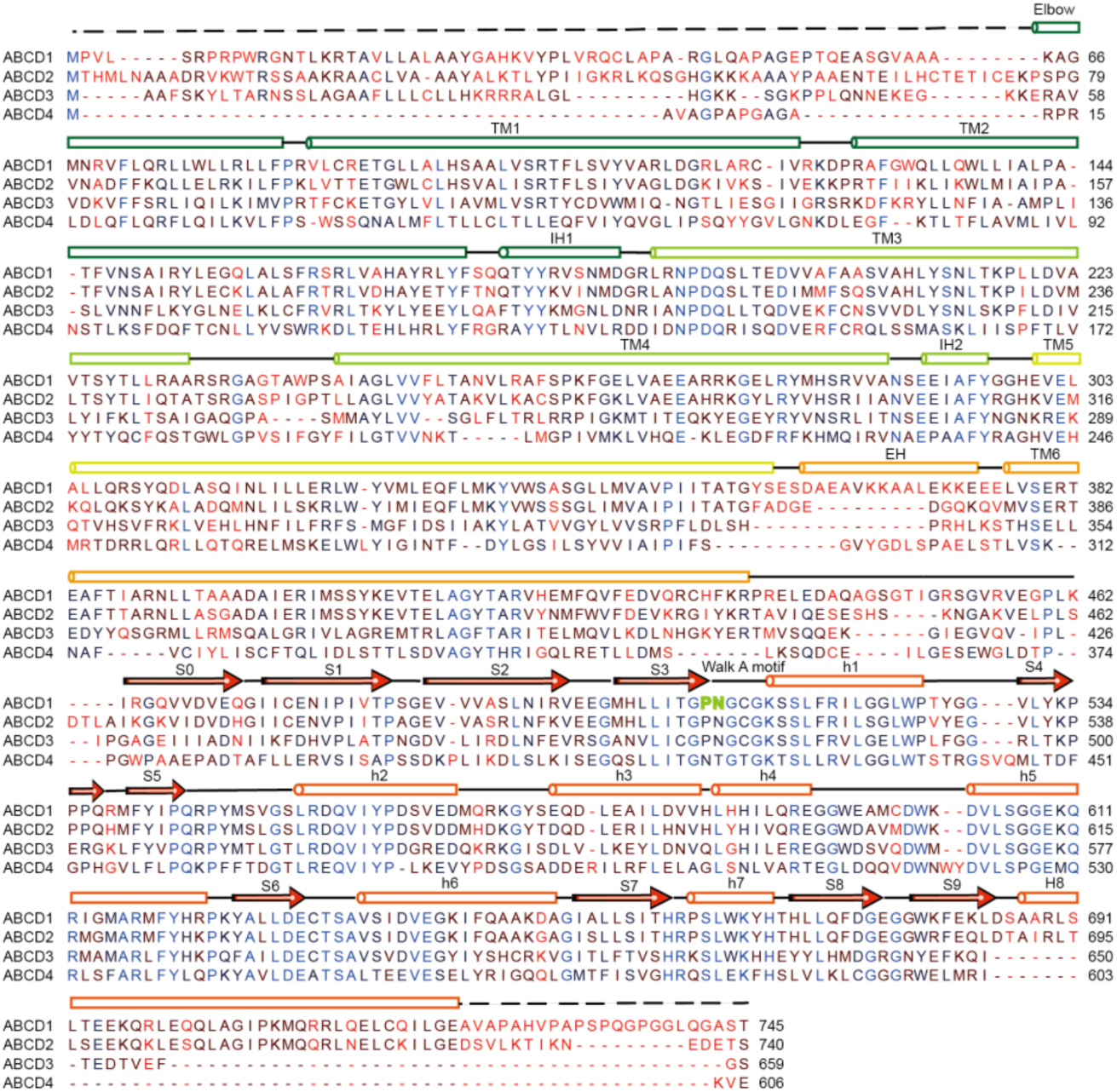
Sequence alignment of human ABCD1 and other members of human ABCD family transporters. Secondary structure assignments are based on the human ABCD1 structure.

**Figure S2.**
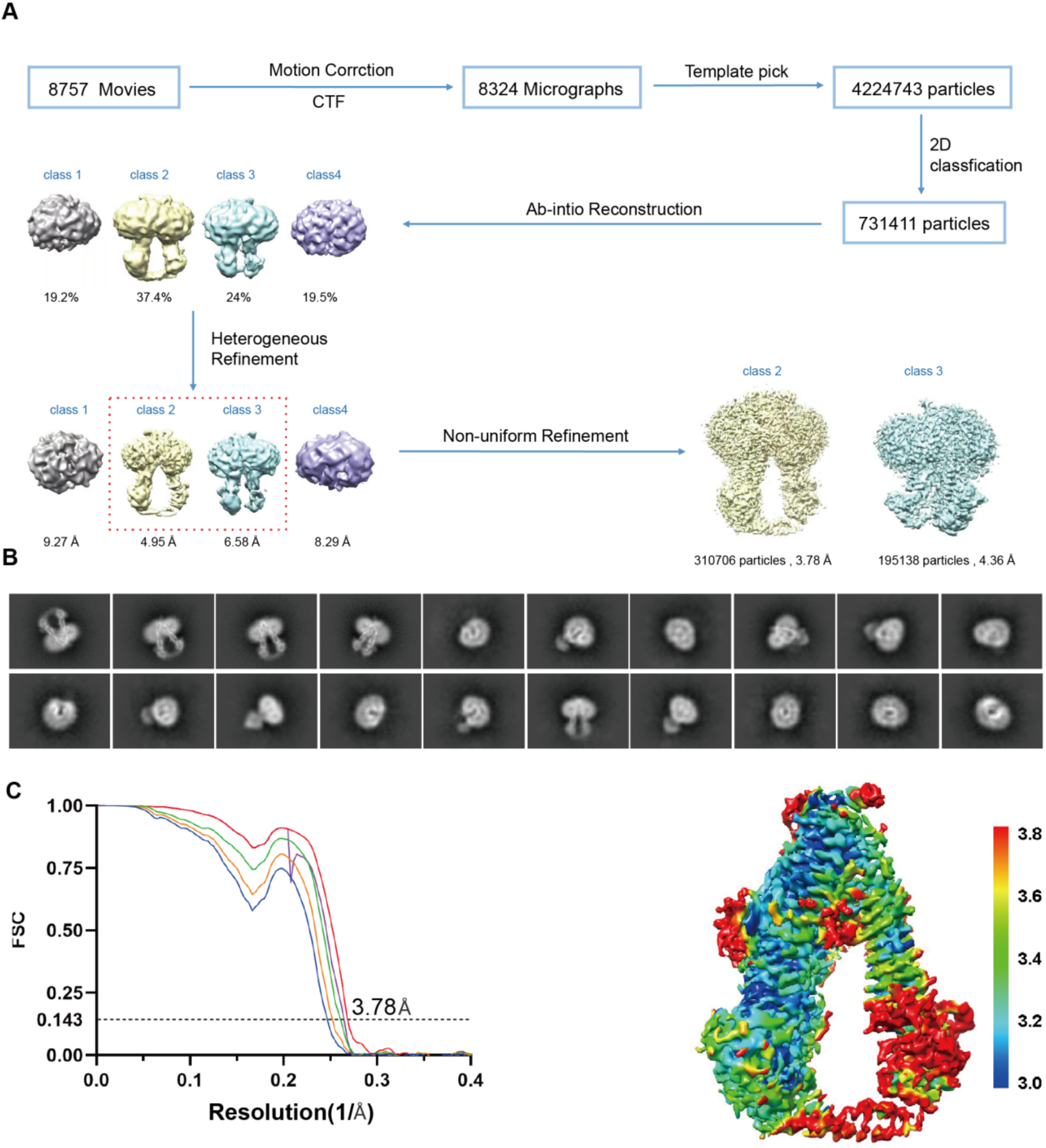
Structure determination of ABCD1-WT-C26:0 complex. (A) Flowchart of image processing for ABCD1-WT-C26:0 particles. (B) Representative 2D classes ofABCD1-WT-C26:0 complex. (C) Gold-standard FSC curves of the final 3D reconstructions and the density map colored by local resolution.

**Figure S3.**
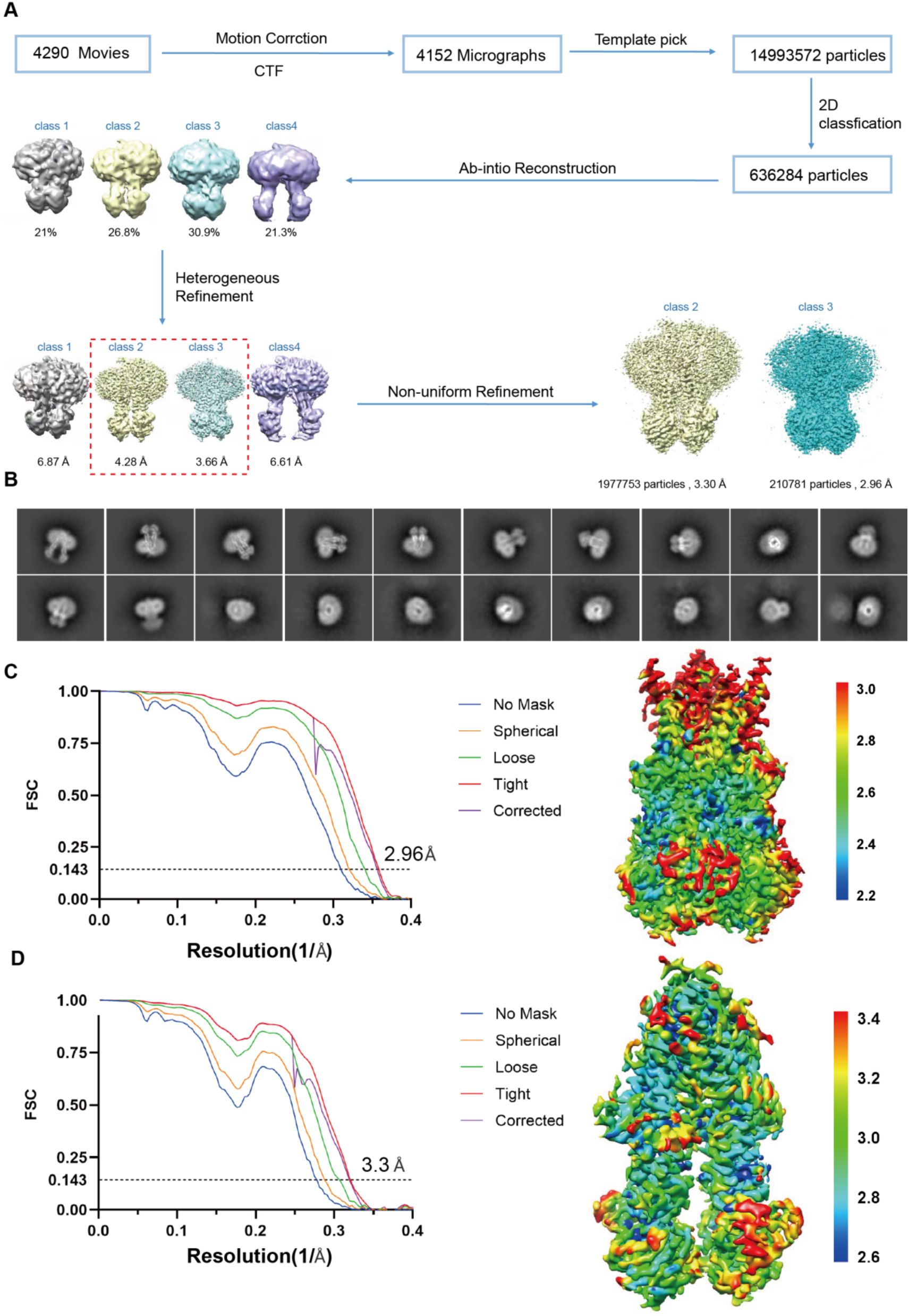
Structure determination of ABCD1-E630Q-ATP complex. (A) Flowchart of image processing for ABCD1-E630Q-ATP particles. (B) Representative 2D classes of ABCD1-E630Q-ATP complex. (C) and (D) Gold-standard FSC curves of the final 3D reconstructions and the density map colored by local resolution for the ABCD1-E630Q-ATP structure in outward facing state and the inward facing state 3 respectively.

**Figure S4.**
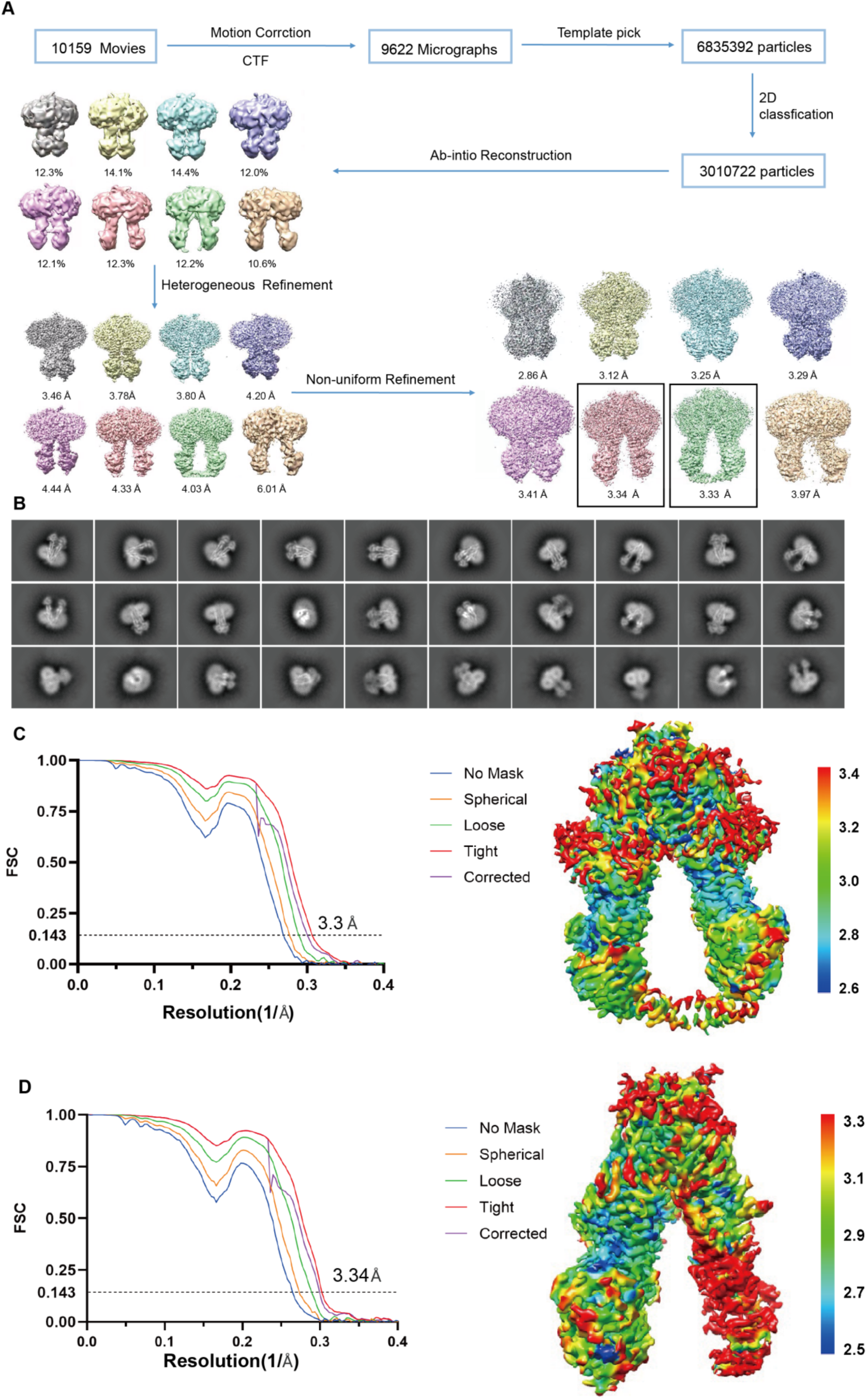
Structure determination of ABCD1-E630Q in the presence of C26:0-CoA and ATP. (A) Flowchart of image processing for the dataset collected on ABCD1-E630Q in the presence of C26:0-CoA and ATP.(B) Representative 2D classes. (C) Gold-standard FSC curves of the final 3D reconstructions and the density map colored by local resolution for ABCD1-E630Q-C26:0-CoA complex in the inward facing state 1. (D) Gold-standard FSC curves of the final 3D reconstructions and the density map colored by local resolution for ABCD1-E630Q-ATP complex in the inward facing state 2.

**Figure S5.**
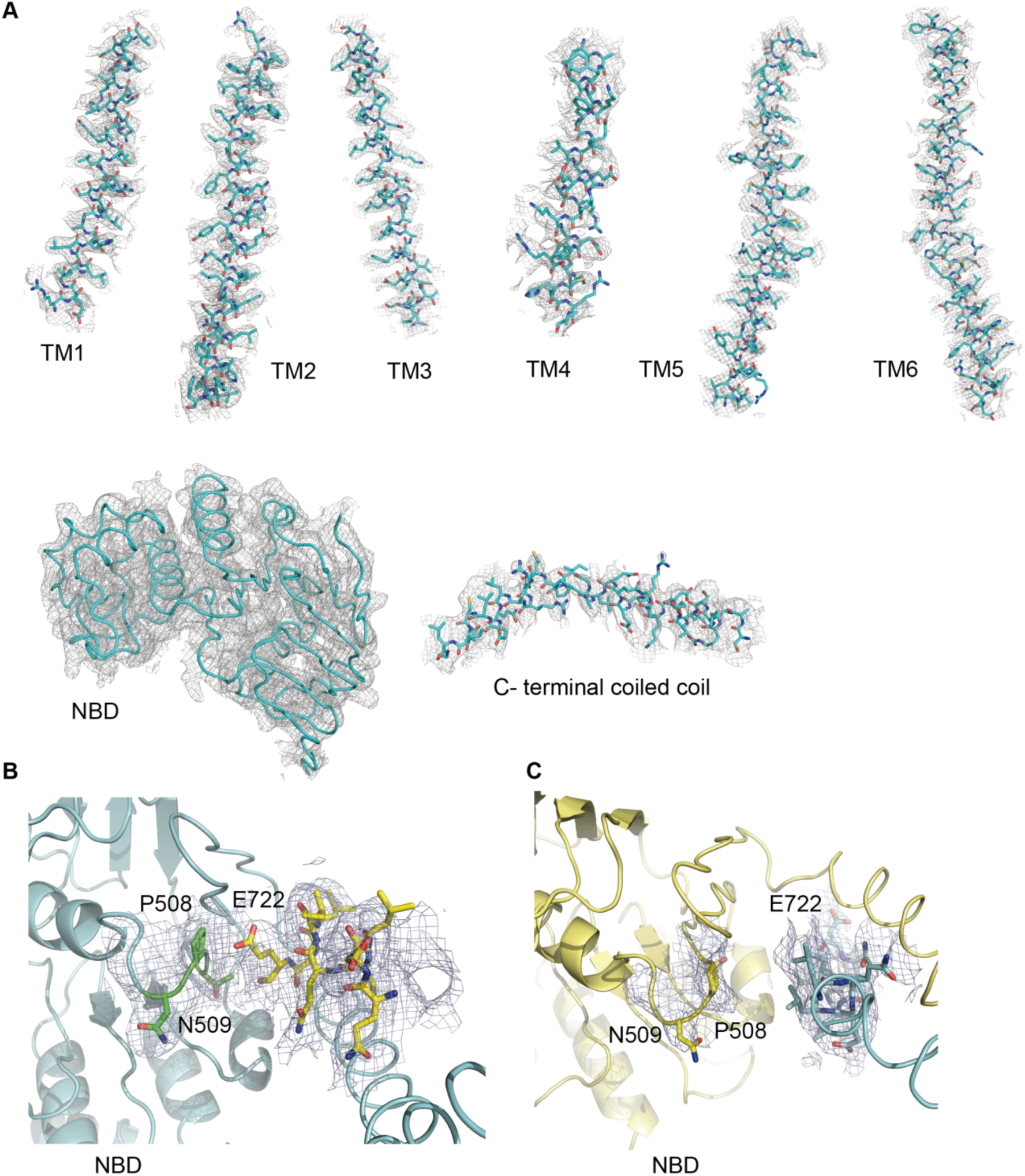
Quality of the EM density, related to Figure 2. (A) EM density of each transmembrane helix, the NBD, and the C-terminal coil. (B) Local structure of the C-terminal coiled coil of one NBD interacting with the Walker A motif of the other NBD. (C) Local structure of the C-terminal coiled coil of one subunit rotating away from the Walker A motif of the other NBD.

**Figure S6.**
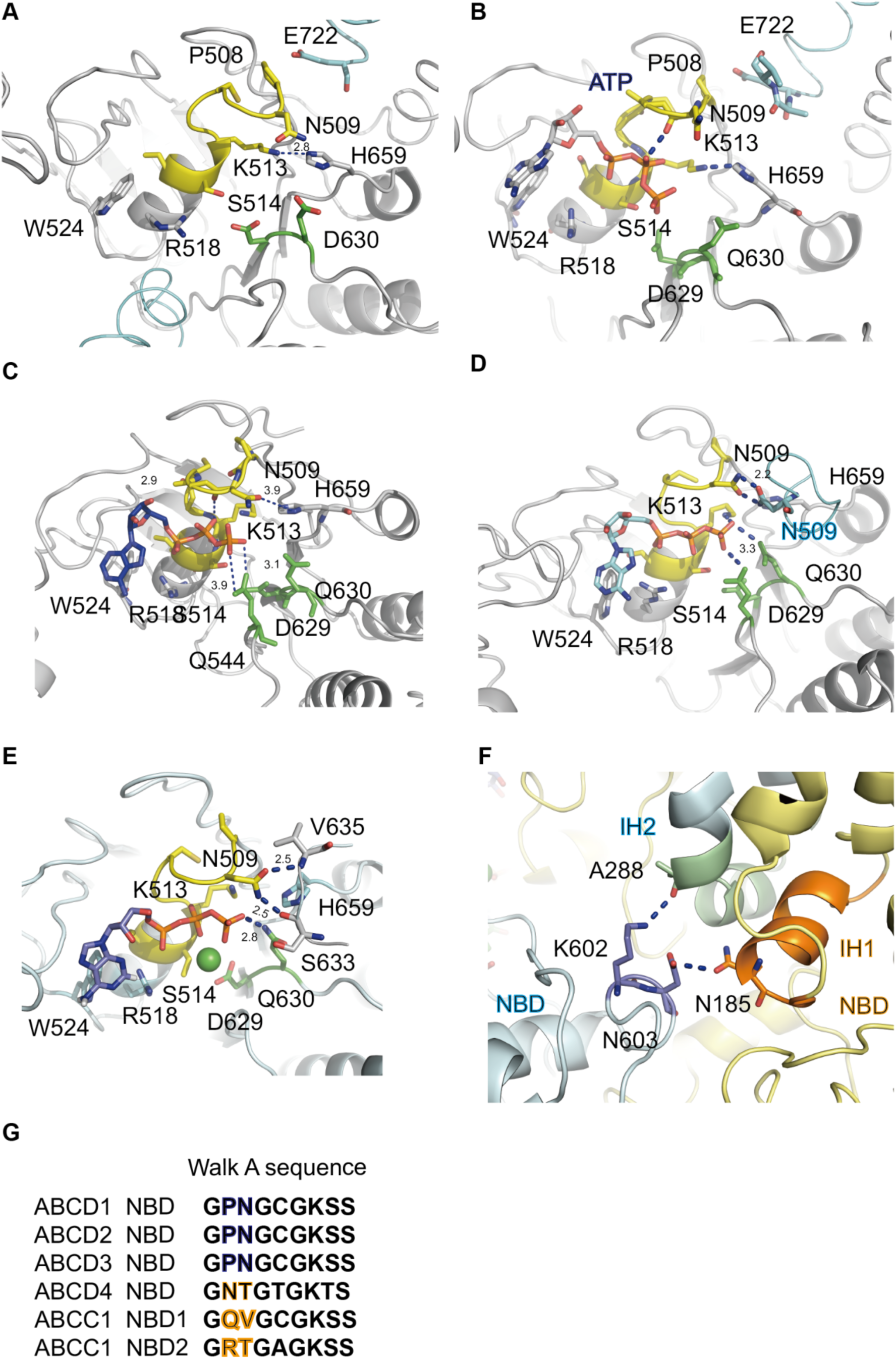
Structural comparison of the ATP binging Site of ABCD1 in different conformational states. (A) and (B) Zoomed in view of the ATP binding site of one NBD for ABCD1-WT-C26:0 complex, and ABCD1-E630Q-C26:0 complex. (C) and (D) Zoomed in view of the ATP binding site of the ABCD1-E630Q-ATP in the inward facing state 2 and state 3 respectively. (E) An ATP molecule interacts with the Walker A motif and the signature motif of ABCD1-E630Q-ATP complex in the outward facing state. (F) Zoomed in view of the NBD-TMD interactions. (G) Sequence comparison of the Walker A motif between human ABCD1 and other ABC transporters.

**Figure S7.**
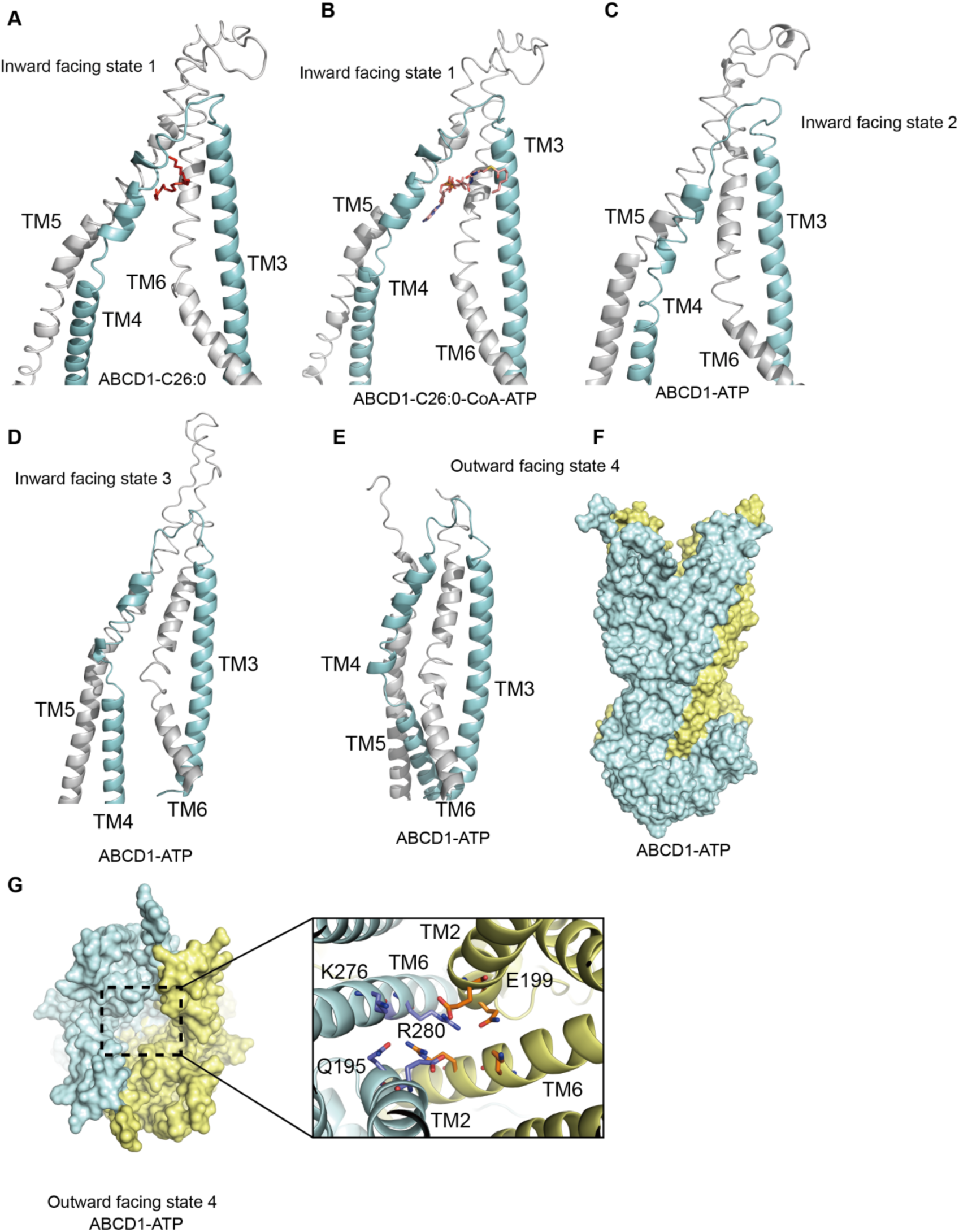
Structural comparison of the 4-helix segments between different conformational states. (A), (B), (C), (D) and (E) The relative positions of 4-helix-TM4, TM3, TM5, TM6 of ABCD1 in different states. (F) Surface representation of ABCD1-ATP in the outward facing state. (G) Charged residues from TM6 and TM2 locates at the bottom of the vestibule opens to the peroxisome lumen.

